# Evolution of self-sustained circadian rhythms requires seasonal change of daylight

**DOI:** 10.1101/2022.04.17.488576

**Authors:** Motohide Seki, Hiroshi Ito

**Affiliations:** Faculty of Design, Kyushu Univesity, Fukuoka, 815-8540, Japan; Institute for Asian and Oceanian Studies, Kyushu University, Fukuoka, 819-0395, Japan

**Keywords:** circadian rhythms, evolution, self-sustained, gene regulatory networks

## Abstract

Self-sustained oscillation is a fundamental property of circadian rhythms and has been repeatedly tested since the early days of circadian research, resulting in the discovery of almost all organisms possessing self-sustained circadian oscillations. However, the evolutionary advantage of self-sustainability has been only speculatively discussed. In this theoretical study, we sought the environmental constraints and selection pressure that drive the acquisition or degeneration of self-sustainability through the process of evolution. We considered a random gene regulatory network dynamics under light cycles and optimized the network structure using an evolutionary algorithm. By designing the fitness function in the evolutionary algorithm, we investigated the environmental conditions that led to the evolution of the self-sustained oscillators. Then, we found that (i) networks showing self-sustained oscillation under constant light conditions are much rarer than those showing damped oscillation and hourglass-type behaviour, and (ii) among several types of fitness-based optimization, networks with self-sustainability property have a markedly high fitness score, especially when we assume that a network has to generate a constantly periodic expression profile regardless of day length. This study was the first to show that seasonality facilitated the evolution of the self-sustained circadian clock, which was consistent with empirical records.

## 1 Introduction

Circadian rhythms are repetitive physiological phenomena with a period of 24 hours. Three unique properties of circadian rhythms—self-sustained oscillations, temperature compensation of the oscillation period, and entrainment by diurnal light or temperature cycles—are shared across kingdoms [1]. Self-sustained oscillation, that is, maintaining the oscillation amplitude without any periodic stimulations, is generally regarded as a more fundamental property than the other two. Temperature compensation assumes self-sustainability because it claims that the period of self-sustained circadian rhythm can be constant under different temperatures. The theory of entrainment argues that external cycles can entrain self-sustained oscillations as long as the applied force is sufficiently strong to shift the phase of the oscillator. Thus, these two properties postulate that the circadian clock system possesses self-sustainability.

Self-sustainability in the circadian rhythms of model organisms has been repeatedly tested since the early days of circadian research. For example, the circadian rhythms of leaf movement last for at least one week under dark conditions, where the environmental time cues are completely shut out [2]. Rodents exhibit circadian rhythms in locomotor activity, a physiologically fundamental measure of circadian rhythm, for several months under constant darkness [3]. However, a few studies have reported two groups of organisms lacking self-sustained circadian oscillations, yeast [4], aphids [5], and purple bacteria [6] are examples of the first group. These organisms possess damped circadian oscillators, indicating that their oscillation amplitudes diminish after being released under constant conditions. Cyanobacteria *Prochlorococcus* and *Hydra vulgaris* exhibited another type of loss of self-sustainability. After these organisms are transferred to constant conditions, the expression levels do not exhibit any peaks and promptly reach an equilibrium state. However, these organisms can sense environmental light and respond to light-dark diurnal cycles [7, 8]. Hereafter, we precisely distinguish the systems that lose self-sustainability using “damped oscillator” for the system that exhibits damping oscillations under constant conditions, such as former examples and “hourglass” for the just-photosensitive system like the latter cases. This classification theoretically corresponds to a stable spiral and a stable node in the trajectory around a fixed point in nonlinear dynamics [9].

Most daily rhythms that have ever been reported satisfy the criteria of self-sustainability, and thus, they are regarded as bona fide circadian oscillations. However, even a damped oscillator and hourglass can show forced oscillations under light-dark cycles, as long as the light signal can affect these semi-clock systems. Thus, regardless of the self-sustainability of the system, the systems behave similarly in a diurnally cyclic environment. This fact brings the question: why has self-sustainability evolved under periodic daily conditions on Earth? Cyanobacteria possess three clock genes *kaiA, kaiB*, and *kaiC* [10], and these clock genes were acquired in the order of *kaiC, kaiB*, and *kaiA* [11]. In addition, the null mutant of *kaiA*, that is, the strain with only *kaiB* and *kaiC*, shows damped oscillation [12], suggesting that the cyanobacterial circadian clock evolved from a damped oscillator to a self-sustained oscillator.

The advantage of self-sustainability has been only speculatively discussed since the early days of circadian research [13]. One can say that developing an endogenous self-sustained oscillator allows the organism to anticipate and prepare daily events to react properly. Self-sustained oscillators maintain their phase as an indicator of their internal state. If circadian machinery has a system that outputs a signal depending on the phase of its clock, the system can measure the time elapsed from the onset or offset of light signals [14]. However, damped oscillators and hourglasses also can exploit this advantage. An hourglass contains system variables that converge to an equilibrium state. Suppose an hourglass system connects with the output system that responds to decay of a variable in hourglass. In that case, the hourglass-type system measures the elapsed time from the switching of environmental light intensity. In principle, a damped oscillator can take advantage of both the phase and decay information if the system can separate these values.

Another hypothesis is seasonality. If the system wants to know the exact time in a day, it is not sufficient to measure the elapsed time from the change in light intensity because the time of sunrise and sunset depends on the season. Thus, a season-independent clock requires additional machinery to absorb the variation in day length. Self-sustained oscillators naturally incorporate an absorber because the entrained phase of the circadian clock depends on the day length; that is, a well-tuned phase response to light stimulation enables daylength-free behaviour. However, a damped oscillator or hourglass-type system can infer the time of day if the system incorporates machinery that preprocesses light signals and then absorbs seasonal variation of photoperiod. Thus, the advantages of the self-sustainability of the circadian clock remain unknown.

This study investigated the environmental constraints and selection pressures that drive the acquisition or degeneration of self-sustainability through evolution. By designing the fitness function of the evolutionary algorithm that leads to self-sustained oscillations, we investigated the environmental conditions under which a self-sustained oscillator is more advantageous.

Several theoretical studies have attempted to explain the evolution of the molecular network of circadian rhythms. For example, Roenneberg and Merrrow [15] considered a coupled system comprising five feedback loops. Each feedback system exhibits damped oscillations under constant conditions; on the other hand, it does forced oscillations under light-dark cycles. When the strength of the coupling between the feedback loops increases, the system shows self-sustained oscillation, even under constant conditions, suggesting that this type of alteration in parameters occurred during the evolution of the circadian clock. Troein et al. [16] used an evolutionary algorithm to search for gene regulatory networks (GRNs) in which gene expression is enhanced just after dawn and just before dusk. They concluded that more complex GRNs were better adapted to seasonal changes in the photoperiod or unpredictable noisy environments. From not the evolution of circadian rhythms but a purely theoretical context, Kobayashi et al. [17] proposed an evolutionary algorithm to find oscillatory GRNs with a specific period length. They also showed that the oscillation period could be easily tuned by modifying the regulations in the optimized GRN.

In this study, we employed a modified version of Kobayashi’s model and used an evolutionary algorithm to examine the evolutionary scenarios of self-sustained oscillations. In particular, considering that there are rarely pure constant light and dark conditions on Earth, we designed a fitness based on the dynamics under light and dark (LD) cycles; that is, we did not explicitly consider self-sustainability under constant conditions for the calculation of fitness, but searched for the conditions in which a self-sustained oscillator behaves more properly than a damped oscillator and hourglass. Through this evolutionary computational search, we tackled on the old question in chronobiology: what drives the evolution of self-sustained oscilation?

## 2 Methods

### 2.1 The model for gene regulatory network

We extended the model of Kobayashi et al. [17] by incorporating an input pathway for the light signal (figure 1*a*). The original model considers a gene regulatory network (GRN) consisting of *N* genes (labelled as gene 1, gene 2, …, and gene *N*) and *M* inhibitory links (*M ≤ N* ^2^). Although activation links were not incorporated into the model, an indirect activation relationship was realized by a chain of two inhibitory links. In addition, each gene is allowed to have a link with itself, namely, direct autoinhibition. Every GRN is represented by a corresponding binary matrix ***A*** = (*a*_*ij*_), where *a*_*ij*_ = 1 if the expression of gene *i* is inhibited by gene *j* and *a*_*ij*_ = 0 otherwise. Note that ***A*** can represent a complex of two or more independent GRNs. We identified such a subdivisible matrix (Appendix A) and excluded it from the analyses below.

**Figure 1.**
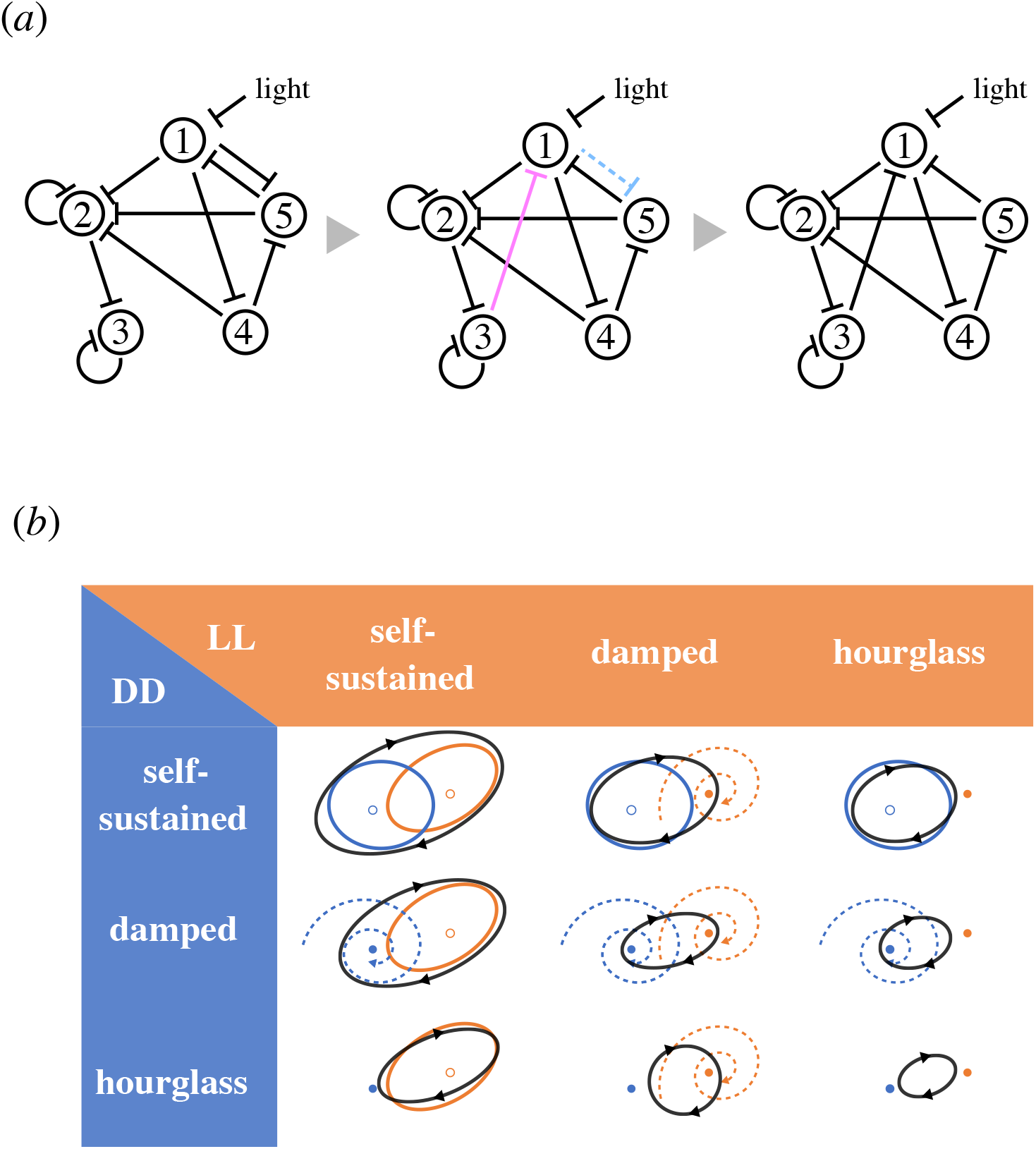
Procedure of evolutionary calculation and classification of systems. (*a*) Illustration of evolutionary calculation. The parent (the left panel) has five nodes and ten inhibitory relationships in this example. One randomly chosen inhibitory relationship (coloured cyan in the middle panel) was removed and one inhibitory relationship (coloured in magenta) was randomly added. Thus, the offspring (the right panel) had the same number of inhibitory relationships as the parents. (*b*) Classification of systems based on the dynamics under light and dark cycles. Systems are classified according to their behaviour under constant darkness (DD, blue) or constant light exposure (LL, orange) into three categories: self-sustained oscillators, damped oscillators, and hourglasses. The three-by-three classes are possible under light and dark cycles, where the system approaches a closed orbit (black), different from the orbits under LL or DD.

In the present study, we analysed a case in which the gene expression 1 was inhibited by light. The light signal was defined as a square wave function of time *t* toggling between 0 (dark) and 1 (light), which was approximated by a continuous function *L*(*t*) in numerical computations to avoid computational errors (see Appendix B). Note that we can consider the case in which the gene expression 1 is inhibited under dark conditions by simply changing the interpretations of *L*(*t*) = 0 and *L*(*t*) = 1.

Expression level of gene *i* at time *t* is denoted by *U*_*i*_(*t*). Dynamics of gene expression is governed by the following ordinary differential equations:

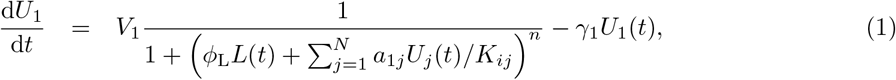

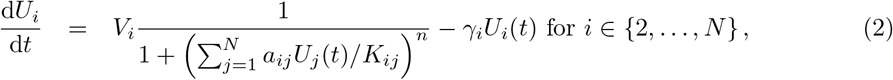

where the first and second terms on the right-hand sides correspond to the production and decay of the gene *i*, respectively. Parameter *V*_*i*_ is the basic production rate (i.e. the production rate without any inhibition) of gene *i*, 1*/K*_*ij*_ is the magnitude of inhibition of gene *i* expression by *j, ϕ*_L_ is the magnitude of inhibition of gene 1 by light, and *γ*_*i*_ is decay rate of gene *i*. The number of parameters was reduced by the nondimensionalized setting *u*_*i*_(*t*) = *γ*_*i*_*U*_*i*_(*t*)*/V*_*i*_ and *ϕ*_*ij*_ = *V*_*j*_*/* (*γ*_*i*_*K*_*ij*_) [17]. Here, we focused on the evolution of the network structure and did not consider the variation in parameters. Thus, we set *ϕ*_*ij*_ = *ϕ*_L_ = *ϕ* for all *i* and *j*, meaning that the strengths of all regulations are equal. Note that gene *j* has the same impact as light on the expression of gene 1, when *a*_*ij*_ = 1 and *u*_*j*_(*t*) = 1. The final form of the system is described as

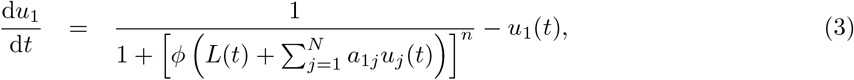

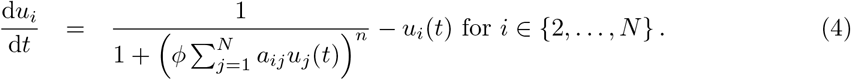

To grasp the basic properties of this system, suppose that there is a simple GRN that has no inhibitory relationships (*M* = 0 and thus *a*_*ij*_ = 0 for all *i* and *j*), although it is an example of a subdivisible network. Under constant darkness, *L*(*t*) in equation 3 can be replaced with zero and thus d*u*_*i*_*/*d*t* = 1 *− u*_*i*_(*t*) for all *i*. We find a uniquely globally stable stationary state 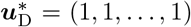, in which every gene shows a constant expression level of 1. Similarly, under constant light exposure, there is a globally stable stationary state 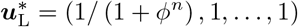 in which the expression of gene 1 is suppressed to 1*/* (1 + *ϕ*^*n*^). Under any given periodic light and dark cycle, the expression level of gene 1 fluctuated within the range of 1 and 1*/* (1 + *ϕ*^*n*^) with the same period as that of the light-and-dark cycle.

In the following numerical simulations, we took an hour as unit time and substituted *N* = 10, *M* = 20, *ϕ* (= *ϕ*_L_ = *ϕ*_*ij*_) = 100, *n* = 3, and a period of light-and-dark cycle (*τ* in Appendix B) as 24, unless otherwise mentioned. A numerical solution up to *t* = 1248 (52 days) was obtained using Mathematica 11.2 (Wolfram Research Inc.). The initial values were set as *u*_*i*_(0) = 0 for all *i* ∈ {1,…, *N*}.

### 2.2 Classification of gene regulatory networks

We developed the following procedure to classify a GRN into three types: self-sustained oscillator, damped oscillator, and hourglass. The first step was to discriminate between the self-sustained oscillator and the other two types based on convergence to a fixed point. The former and latter should not and should converge to a fixed point after a long time, and thus a standard test for convergence was applied. We measured the distance in the phase space ***u*** = *{u*_1_, *u*_2_,…, *u*_*N*_}, between the states of the system at two time points*t*_*a*_ and *t*_*b*_ by Δ(*t*_*a*_, *t*_*b*_) = max_*i*_ |*u*_*i*_(*t*_*a*_) − *u*_*i*_(*t*_*b*_)|. We classified systems that satisfy

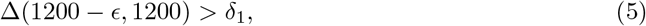

where *ϵ* = 10^−2^ and *δ*_1_ = 10^−6^, as candidates for a self-sustained oscillator. To check the regular periodicity of the candidate and exclude chaotic behavior, we carefully examined the oscillation period *T*, which satisfies *u*(*t*) = *u*(*t* + *T*). The gene expression profiles that showed most frequently peaks from the 41th day to the 50th day was chosen among the *N* genes. Suppose that we found *n*_*p*_ peaks for the profile and denote times of those peaks as 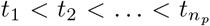. We assumed that the system at time *t*_*k*_ remained in the vicinity of 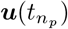 if *t*_*k*_ satisfied

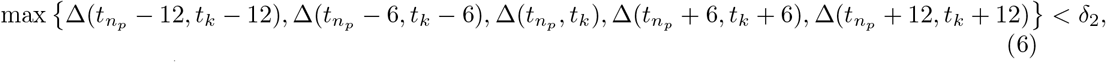

where *δ*_2_ = 10^−4^. Denoting the greatest peak time among 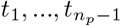 that satisfies equation 6 as 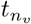, we obtained the oscillation period as 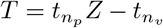. The difference between *t*_*Z*_ and the first-found *t*_*k*_ that satisfies equation 6, *t*_*Z*_ − *t*_*k*_ was considered as the period. If none of the peaks satisfies equation 6, we regarded the period of oscillation as “more than 10 days or chaotic.”

The second step for a system classified as a non-self-sustained oscillator is to discriminate between whether it was a damped oscillator or an hourglass. We regard ***û*** = (*u*_1_(1200), *u*_2_(1200),…, *u*_*N*_ (1200)) as a stable fixed point, and numerically obtained the leading eigenvalue of Jacobian matrix around the fixed point. The system is classified as a damped oscillator if the leading eigenvalue is a complex number. and hourglass otherwise (see [9]). A similar procedure was applied to a GRN under light-dark cycles. The GRN under this condition is classified as a driven oscillator if equation 5 was satisfied. For driven oscillators, equation 6 was applied to estimate the period length of the driven oscillation.

### 2.3 Optimization using evolutionary algorithm

We applied an evolutionary algorithm to obtain a 10-gene-20-regulation network in which gene 10 shows an expression profile close to a given target profile (see each section). A cost value was numerically calculated using the following temporal integration:

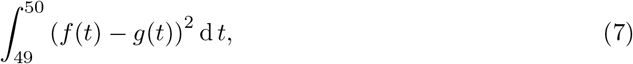

where *f*(*t*) and *g*(*t*) are the target and actual profiles of gene 10, respectively. One generation consists of mutation and selection. In the mutation process, we generated an offspring GRN by copying the parental GRN, deleting a randomly-chosen one of the *M* inhibitory relationships, and then added a new inhibitory regulation chosen from the *N* ^2^ − *M* candidates (figure 1*a*). In the selection process, either the parental GRN or offspring GRN is selected as the parent in the next generation, following the rule below. The offspring is selected as the next parent if (i) it is a connected GRN (see Appendix A), and (ii) it shows the same or smaller value of the cost function than that of the parent. Otherwise, the current parental GRN is selected as the next parent. For each set of a target profile and an environment, we ran 1,000 independent trials with different initial parental GRNs and lasting 2,000 generations.

### 2.4 Classification of networks based on bifurcation

We varied the value of the parameter *ϕ* in the evolutionarily optimized self-sustained oscillators to detect a supercritical Hopf bifurcation. We regarded that a system showed a supercritical Hopf bifurcation when the oscillation amplitude at steady-state continuously decreased to zero and period length did not change significantly as we reduced the value of *ϕ*.

## 3 Results

### 3.1 Self-sustained oscillation is rarer than damped oscillation and hourglass

We first confirmed that the self-sustained oscillators were rarer than the other two types. Specifically, we randomly generated gene regulatory networks (GRNs) consisting of 10 genes and 20 intergenic relationships (i.e. *N* = 10 and *M* = 20) to obtain 1,000,000 connected GRNs. To complete the process above, we generated 1,131,417 GRNs because subdivisible GRNs were generated 131,417 times. We also found that very few (123) GRNs were generated twice, and no GRNs were generated more than twice. Only 6.4% of the GRNs exhibited self-sustained oscillations under either constant light (LL) or constant dark (DD) conditions (figure 2*a*). GRNs showing damped oscillation under at least one condition were found much more frequently (44.2%). The majority of GRNs (51.7%) were hourglasses under both constant conditions. GRNs that showed self-sustained oscillation under DD were slightly more frequent than those under LL. This asymmetry between LL and DD could be due to our assumption that the expression level of gene 1 is kept near zero under LL. Effectively, this reduces the number of network member genes (*N*) from 10 to 9 (see below and figure 2*b* for the effect of *N*). In addition, the diagonal elements in figure 2*a* have greater values than expected under the assumption that behaviours under LL and DD were completely independent.

**Figure 2.**
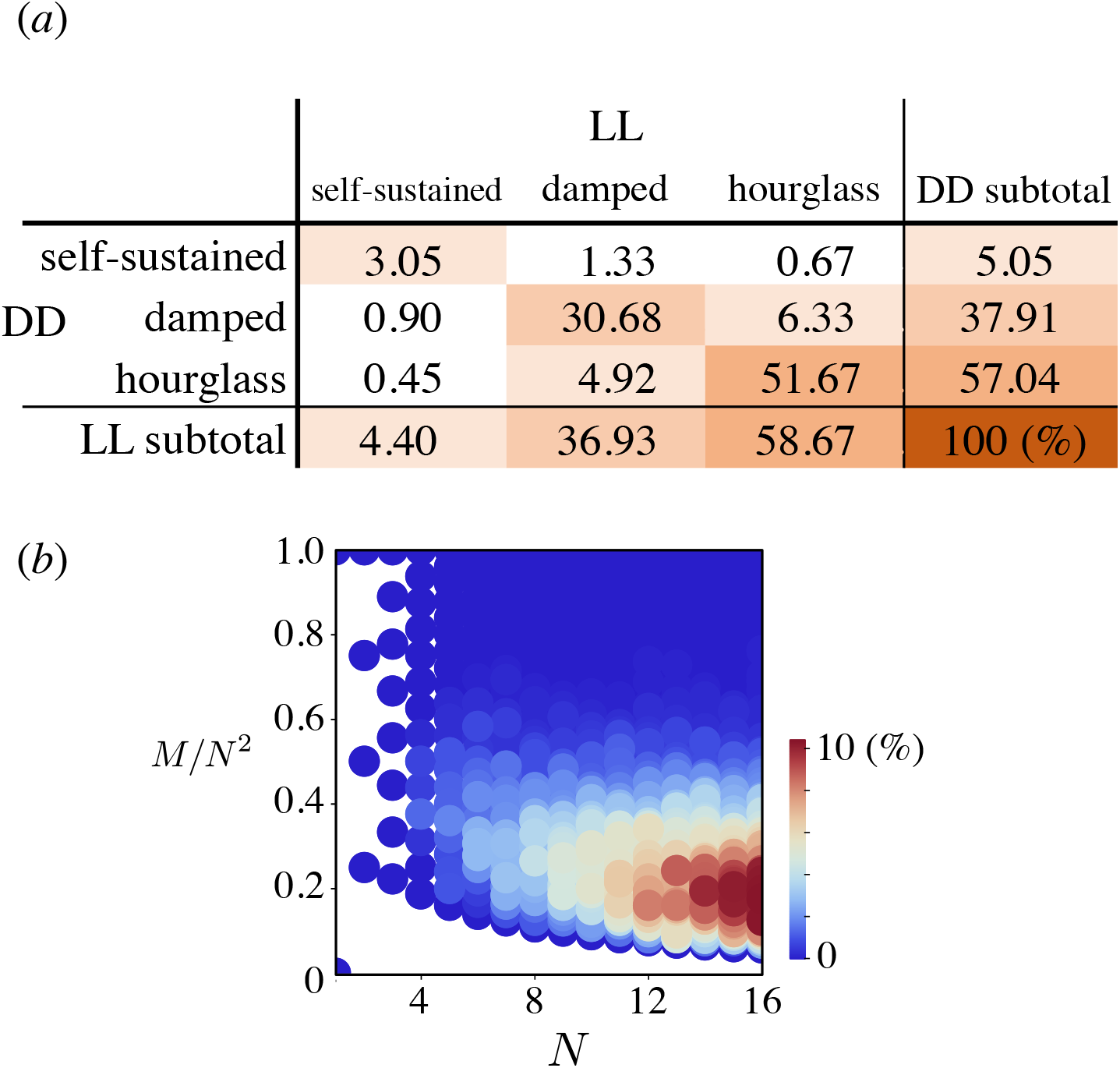
Dynamics of the randomly generated gene regulatory networks (GRNs). (*a*) Frequencies of the classes in one million connected GRNs consisting of 10 genes and 20 inhibitory relationships (i.e., *N* = 10 and *M* = 20). (*b*) Heat map indicating frequencies of self-sustained GRNs plotted as the number of genes (*N*) versus the filling rate of inhibitory relationships (*M/N* ^2^). Note that a *N*-gene network can have at most *N* ^2^ inhibitory relationships, including self-to-self inhibitions. For each pair of *N* (∈ {2,…, 16}) and *M* (∈ {1,…, *N* ^2^})), we test all possible GRNs if the number of possible GRNs (*N* ^2^)!*/*[(*N* ^2^ − *M*)! · *M*!] was less than 1,000, or 1,000 randomly generated GRNs otherwise.

We then simulated a 12 h light/12 h dark environment (hereafter denoted as 12L12D) and confirmed that all GRNs exhibited oscillations with the period of 24 h. With such a setting, the gene expression dynamics under dark and light conditions are governed by different systems with different values (0 in the dark and 1 in the light) for parameter *L*(*t*), and thus showed different sets of fixed points and even different oscillatory modes (figures 1*b* and S1). Even a GRN behaving as an hourglass under both DD and LL conditions behaved as a daily oscillator because gene expression dynamics were periodically switched between two stable fixed points under dark and light conditions. Note that the numbers of fixed points under the LL and DD are not generally different, i.e., a fixed point under LL often has a correspondent one under DD. However, the stabilities of the fixed points differ due to Hopf bifurcation or transcritical bifurcation when the bifurcation point for parameter *L* exists between zero and one [9].

In addition, we examined the dependency on the number of genes *N* and on the number of edges *M*. The greater *N* and *M*, that is, a larger scale or a higher complexity of a GRN, favours self-sustained oscillation (figure 2*b*). Strong gene-to-gene suppression (i.e. greater *ϕ*) also facilitates self-sustained oscillations (figure S2*a*). A self-sustained oscillator involving more genes and/or a moderate number of edges had a longer period on average (figure S2*b*).

### 3.2 Seasonal environment can favour self-sustainability

To specify selective force(s) that can lead to a nonrandom evolution of self-sustained oscillation, we performed our evolutionary algorithm and collected the 10-genes networks under some LD conditions in which the expression of gene 10 had a target profile. We specified environmental conditions that facilitated the self-sustainability by observing the type of the GRNs obtained through the evolutionary algorithm. The best GRN in an environment was defined as the GRN with the smallest cost value among outcomes of 1,000 trials of the evolutionary computation.

First, assuming a 24-hour periodic sinusoidal curve with a peak at dawn as the target profile, we performed evolutionary computation under the 12L12D environment. A majority of the final outcomes showed self-sustained oscillation neither under dark nor under light (figure 3*a*; the top panel of figure 3*e*). Using a sinusoid curve with a peak at dusk resulted in a similar distribution. (figure 3*c*).

**Figure 3.**
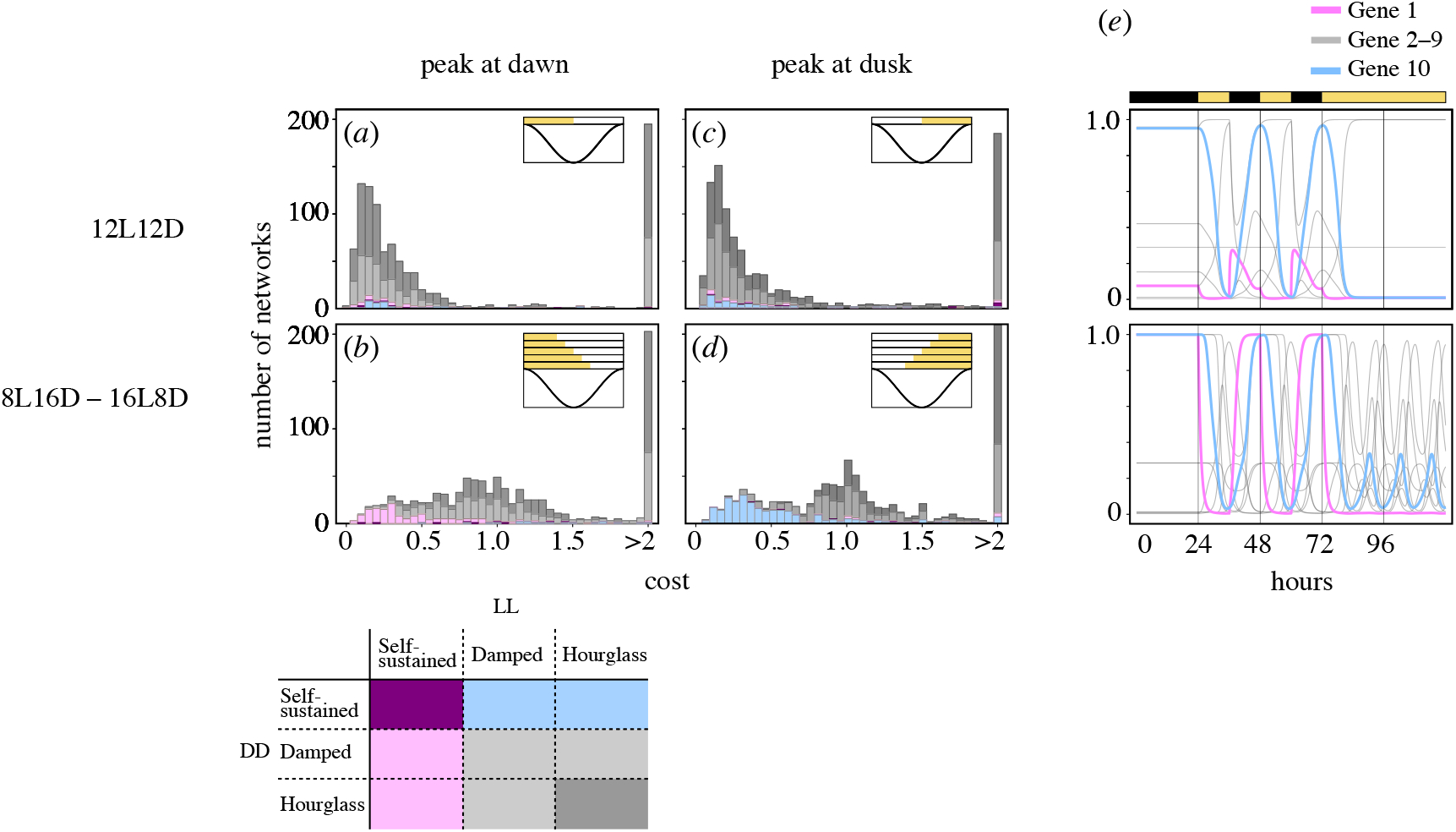
Results of evolutionary calculations in which a sinusoidal curve with a peak at dawn or dusk was assumed to be the target profile for the gene 10 expressions. (*a*–*d*) Distribution of cost values for the outcome gene regulatory networks (GRNs) from 1,000 different trials with different initial GRNs. The horizontal axes represent cost values, and the vertical axes indicate frequencies of GRNs. A cost value represents the deviation from the target profile. Histograms arranged in the left and right columns show outcomes of simulations in which the target expression profiles for gene 10 were sinusoidal curves with peaks at dawn and dusk, respectively. In conditions labelled as 12L12D (*a, c*), a GRN was repeatedly exposed to a 12-hour-light and 12-hour-dark cycle 50 times, and a cost value was calculated on the last 24 hours. In conditions labelled as 8L16D–16L8D (*b, d*), a GRN was independently exposed to 8L16D, 10L14D, 12L12D, 14L10D, and 16L8D environments, and cost values of those five environments were averaged. GRNs were classified according to behaviours under constant darkness and constant light exposure: purple for a GRN showing self-sustained oscillations under both constant environments, magenta and cyan for a GRN showing self-sustained oscillations only under light and dark conditions, respectively, dark grey if it does not show self-sustained or damped oscillations in any constant environment, and light grey otherwise. (*e*) Expression profiles of the genes in the GRNs with the lowest cost value in the 12L12D environment (top panel) and in the 8L16D–16L8D environment (bottom panel) when the target profile has a peak at dawn. Both GRNs were exposed to constant darkness for 24 hours, 12L12D condition for 48 h, and then under constant light. The GRN selected under the 8L16D–16L8D environment showed self-sustained oscillation under constant light.

We next considered a seasonal environment mimicking a temperate zone. Specifically, the cost value of a GRN was defined as the average of five cost values independently calculated under the 8L16D, 10L14D, 12L12D, 14L10D, and 16L8D environments. With this setting, most GRNs with lower cost, that is, higher fitness, behaved as self-sustained oscillators under constant light condition while damped oscillators or hourglasses under constant dark condition (figure 3*b*, the bottom panel of figure 3*e*). In addition, the final outcomes had on average greater cost values than those in the previous computation with the single 12L12D environment (compare figure 3*a, c* with 3*b, d*), indicating the difficulty of achieving similar expression profiles under multiple light environments. Changing the peak time of the curve from dawn to dusk resulted in qualitative alterations in the outcomes; self-sustained oscillators only under dark was frequently observed in place of self-sustained oscillators only under light (compare figure 3*b* and 3*d*).

Of the GRNs optimized through evolutionary computation under seasonal environment, those showing self-sustained oscillations under light (magenta or purple cases in figure 3*b*) were further analysed to determine how they lost their self-sustainability (i.e. which type of bifurcation they followed) when the parameter of the system *ϕ* was reduced from 100. Most (161 of 164) GRNs transitioned from self-sustained oscillators to non-oscillators via supercritical Hopf bifurcation (figure S3*a*), and the other three GRNs appear to show other types of bifurcation (figure S3*b*).

We tested the generality of the above-obtained conclusion that self-sustained and non-self-sustained oscillators have higher fitness than the other types under seasonal and aseasonal environments, respectively. Changing the trough time of the target profile from 12 h to 8 or 16 h after dawn did not qualitatively change this conclusion (figure 4). It was clarified that self-sustained oscillations are especially useful to revert the expression level of gene 10 at midday (figure 4*g*). In addition, we perturbed the wave form of the optimal profile, i.e., although assuming the only sinusoidal curve for the optimal profile so far, we considered the case where further changing the trough time is dependent on daylength. This flexible wave form yielded lower frequencies of self-sustained oscillators among the outcomes compared with the sinusoidal profile (figure S4*a, b*). We also perturbed function form of the target profile by sharpening the peaks and flattening the troughs (the bluish and reddish curves in figure S4*c*, respectively). Such a form is modelled on circadian gating, which realizes a long inactive or insensitive phase often through regulation by the circadian clock [18]. It turned out that self-sustained oscillators can appear even in aseasonal conditions when the gating-type profile is optimal (see figure S4*d*). Regarding seasonal conditions, our evolutionary computation did not detect any GRNs with high fitness (figure S4*e*).

**Figure 4.**
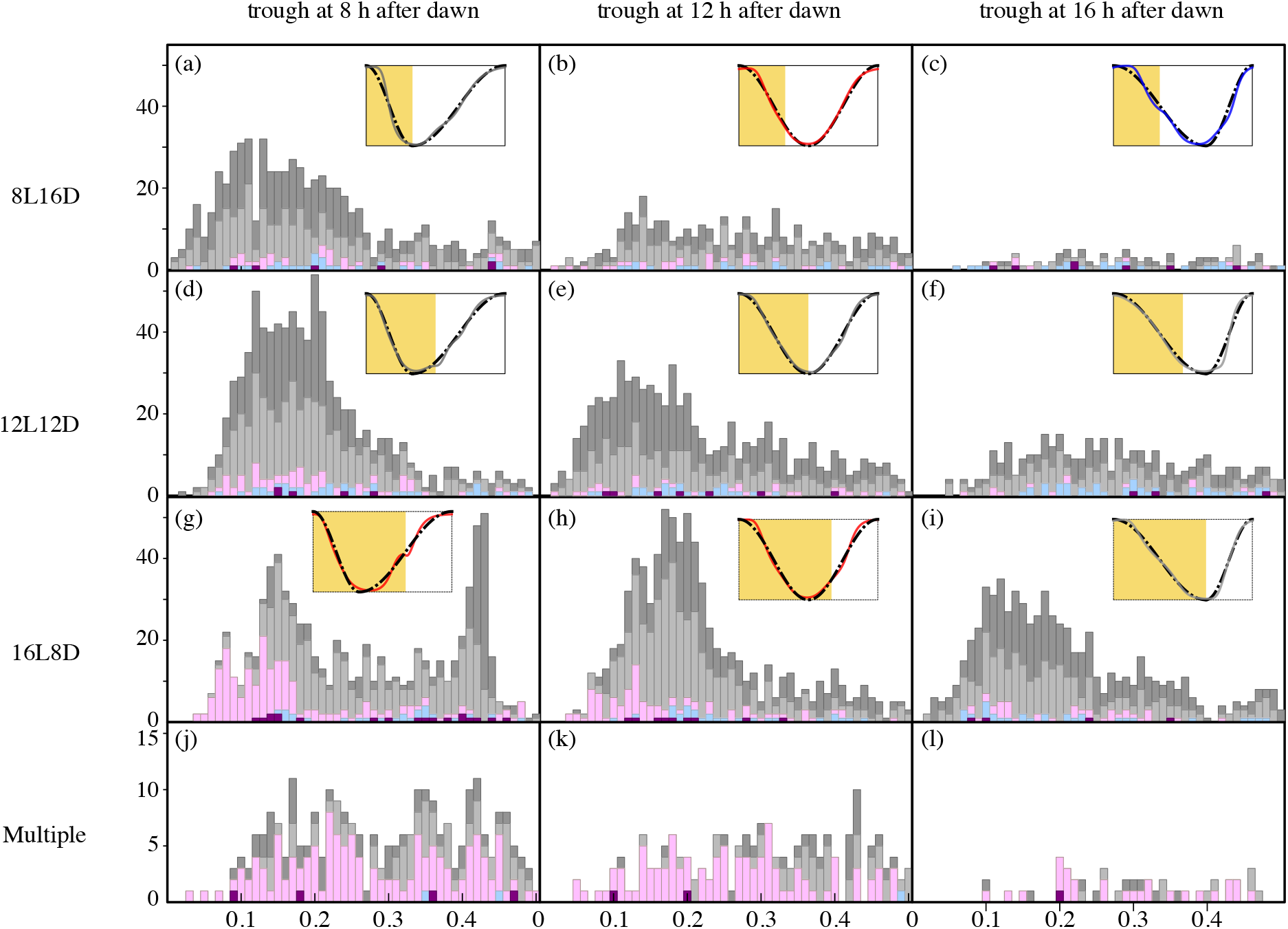
Distribution of cost values for the outcome gene regulatory networks (GRNs) from 1,000 different trials with different initial GRNs under the cycles with different photoperiods. The horizontal axes represent cost values, and the vertical axes indicate frequencies of GRNs. A cost value represents the deviation from the target profile. Histograms arranged in the left, middle, and right columns show outcomes of simulations in which the target profiles were assumed to be sinusoidal-like curves with troughs at 8, 12, and 16 h after dawn, as shown by dotted-line curves in the embedded plots. In the first, second, and third rows, a GRN was exposed to 8-hour-light and 16-hour-dark (8L16D), 12L12D, and 16L8D environments, respectively, repeatedly 50 times. Then a cost value was calculated on the last 24-hour period. In the bottom row, a GRN was independently exposed to 8L16D, 10L14D, 12L12D, 14L10D, and 16L8D environments, and cost values of those five environments were averaged. GRNs were classified according to behaviours under constant darkness and constant light exposure: purple for a GRN showing self-sustained oscillations under both constant environments, magenta and cyan for a GRN showing self-sustained oscillations only under light and dark conditions, respectively, dark grey if it does not show self-sustained or damped oscillations in any constant environment, and light grey otherwise. Coloured curves in the embedded plots are expression profiles of gene 10 in the GRNs with the lowest cost values. Note that panels *e* and *k* are rescaled duplications of figure 3*a* and 3*b*, respectively.

## 4 Discussion

This study examined the evolution of self-sustainability in the circadian clock by observing the behaviour of random gene regulatory networks (GRNs) under diurnal cycles. We found that GRNs showing self-sustained oscillation under a constant light condition are rarer than those losing self-sustainability, that is, those showing damped oscillation or hourglasses. However, GRNs with self-sustainability showed higher fitness than the other types in evolutionary computation when they had to generate an expression profile with a constant form regardless of day length (figure 3*b, d*), whereas a non-self-sustained system was preferable when the systems could utilize the environmental switch, that is, when the target gene expression showed a trough at dusk (figure 3*a, c*). These results suggest that seasonal variation of day length is a driving force of evolutions of self-sustainability.

More precisely, self-sustainability under constant light contributes to the realization of a trough under light period in a cycle (figure 4*a, g* ; 4*b, h*). Moreover, the trough at mid-day was more effective (compare figure 4*g* with 4*h*). These results suggest that self-sustainability in LL conditions can contribute to both peaks and troughs during the light period of LD cycles. In other situations, a damped oscillator or hourglass may work as an alternative to a self-sustained clock. The fact that shifting the peak time to the beginning of the dark period accelerates the preference for self-sustainability in DD supports this hypothesis (see figure 3*d*). Thus, a higher latitude can benefit self-sustainability because seasonal conditions necessarily contain this self-sustained-dominant condition.

We also found that GRNs with the property of self-sustainability were exclusively favoured to generate an identical gene expression profile regardless of the day length (figure 4*i, j, k*), whereas damped oscillators and hourglasses seemed to have difficulty generating a consistent profile independent of day length. The endogenous season-independent timing system is implicitly presumed by the external coincidence model, stating that a seasonal event (e.g. flowering or hibernation) is launched when the photoperiod coincides with the endogenous rhythm [19]. In the example of Arabidopsis flowering, CONSTANS (CO) protein is known to induce floral transition when its amount is above a threshold and to be degraded under dark. In addition, *CO* gene expression is regulated by the circadian clock. Given that the expression profile of *CO* is not greatly affected by daylength and reaches above the threshold at a certain timing, say 16 h, flowering is and is not induced when the day length is longer and shorter than 16 h, respectively [20].

Our finding that seasonality can favour the evolution of self-sustainability is consistent with comparative studies on the variation in circadian rhythms across latitudes. *Synechococcus*, a cyanobacterial genus known to have a circadian self-sustained oscillator, is widely distributed both under seasonal and aseasonal environments. In contrast, *Prochlorococcus*, a genus with a light-driven 24-hour oscillator, are absent in high-latitude regions with strong seasonality [21]. Previous theoretical studies focusing on the circadian machinery in these two genera deduced another explanation concerning noise sensitivity [22, 23], which is not mutually exclusive with our seasonality hypothesis. Using duckweeds species in the genus *Lemna* distributed between tropical and subarctic zones, Isoda et al. [24] found a tendency similar to the above. They examined the self-sustainability of the circadian rhythm of plants in a wide range of temperature conditions and reported that the species inhabiting colder (i.e., higher latitudinal) regions had more stable self-sustainability than those inhabiting lower latitudinal regions, indicating the importance of the self-sustainability in seasonal environments and/or non-necessity of self-sustainability in aseasonal environments. We expect other undescribed examples of organisms lost their self-suitability in tropical areas.

Self-sustained oscillation is more likely to appear in larger-scale GRN involving many members and a moderate number of regulatory relationships (figure 2*b*). This may correspond to the fact that self-sustained circadian clocks are generally large-scale. The conceptual model of the *Drosophila* circadian clock provided by Rivas et al. [25] involved 12 genes. Sanchez and Kay [26] states that “minimal architecture” of the circadian clock in *Arabidopsis thaliana* consists of 10 genes.

The perturbation of the parameters of the system in the optimized GRNs mostly caused supercritical Hopf bifurcation (figure S3), meaning that the oscillation amplitude, rather than the period, depends more on the parameter. The preference for Hopf bifurcation in our algorithm suggests that circadian clocks in nature can be surrounded by a damped oscillator region in the parameter space. Lowering the ambient temperature nullifies cyanobacterial circadian clocks via Hopf bifurcation [27]. This result contrasts with the fact that the period of most GRNs obtained via Kobayashi’s algorithm is sensitive to *ϕ* [17]. This robustness against perturbation at parameter suggests that circadian rhythms are not merely self-sustained but also share other common properties. Further research would hint at the enigma in chronobiology: why all of the circadian clocks reported so far have transcriptional and translational feedback loops [28].

One limitation of the present study is that our research considered GRNs with only nonlinear inhibitory regulations [17]. This model is relatively abstract; however, studies using other models (e.g. the Boolean network model [29]) are required to exclude the model dependency of our results. Another limitation is the use of a simple evolutionary algorithm, in which one gene-to-gene regulatory relationship dissolves and a novel relationship appears. A real biological mutation would alter magnitude of the existing regulation (the value of *ϕ*_*i*_ in the model) or enlarges/shrinks the scale (i.e., the number of genes involved) of the GRN. Thus, a more complex mutation system is required to trace the evolutionary pathway of the circadian clock more precisely. In addition, the use of an evolutionary algorithm could result in relatively few GRNs showing a gene expression profile that is highly similar to a target profile. A more efficient method, such as the Markov chain Monte Carlo method [30], would provide a greater number of good-fit GRNs, which would help to understand selective force on circadian clocks.

There have been few reports on the evolution of self-sustained circadian rhythms. Laboratory evolution assays are straightforward for this topic; however, no reports have succeeded in demonstrating the appearance of the circadian clock in the laboratory. Drawing a phylogenetic tree based on genome sequences can provide a history of circadian molecular machinery, but not the adaptive mechanism. Instead, we have numerically introduced a tractable approach based on an evolutionary algorithm, which provided an unbiased finding that seasonality facilitated the evolution of the self-sustained circadian clock. We expect that confirmation of our idea by experiments and further theoretical studies should advance the study of the evolution of circadian rhythms.

## Appendix A An algebraic method to check subdivisibility of networks

A network is considered connected if it contains at least one one-way path between every pair of nodes. It is called strongly connected if it includes at least one path directed from every node to another. A network with *N* nodes represented by an adjacency matrix ***M*** is strongly connected if (***I*** + ***M***)^*N−*1^ is a positive matrix, where ***I*** is an *N* × *N* identity matrix [31].

To check whether a graph represented by an adjacency matrix ***A*** is connected, we define a new *N* × *N* matrix ***B*** = (*b*_*ij*_), where

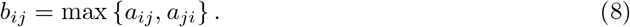

The focal network is connected if the network represented by ***B*** is strongly connected, that is, if the matrix (***I*** + ***B***)^*N−*1^ is positive.

## Appendix B Continuation of the periodic binary function

Because the method used in this study to obtain numerical solutions for differential equations can cause a numerical error in analysing a system involving a sudden or unsmooth change in a variable, it is convenient to use smooth and continuous functions. The master equations describing the proposed model (equation 3 and 4) contain a discontinuous function, *L*(*t*), to represent periodic change in the light status. By revising a function previously proposed by Pokhilko et al. [32], we approximated *L*(*t*) using the following function:

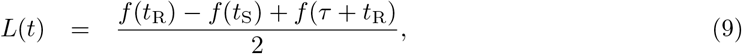

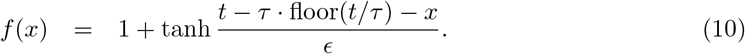

where *t*_R_ and *t*_S_ are the phases of sunrise and sunset, respectively; *τ* is the period of a light-and-dark cycle; and *ϵ* is the duration of the transition between 0 and 1. We have used *τ* = 24 and *ϵ* = 10^−2^.

## Acknowledgements

We thank T. Takada and H. Kato (Hokkaido University) for advice on mathematical methods; N. Kon and Y. Fukada (University of Tokyo), A. Kubota and M. Endo (NAIST) for fruitful discussions.

## Data accessibility

Code implementing the model of gene regulatory network (GRN) and search for the optimal GRN is available on Github https://github.com/hito1979/RSB2022.

## Author’s contribution

M.S. and H.I. conceived the study, carried out the theoretical analysis and numerical study, and wrote the paper. Both authors approved the final manuscript for publication.

## Figure Captions

**Figure S1**

Example of a system with *N* = 2 and *M* = 4 that exhibits hourglass-type dynamics under both constant darkness (DD) and constant light exposure (LL). Horizontal and vertical axes represent expression levels of genes 1 and 2, respectively. (*a, b*) Trajectories under DD (*a*) and LL (*b*). The fixed point locates at different points under LL and DD. (*c*) The system approaches a closed orbit during light and dark (LD) cycles (4L4D in this example). Figures *a* and *b* are superimposed to c.

**Figure S2**

Dependency of properties of a randomly generated gene regulatory network (GRN) on parameters. (*a*) Proportion of GRNs showing self-sustained oscillation under dark versus *ϕ*. (*b*) Median period of self-sustained GRNs under dark versus normalized number of inhibitory relationships. Period zero means that no self-sustained GRNs were generated within 1,000 trials.

**Figure S3**

Dependency of oscillation period and amplitude on the parameter *ϕ*. (a) Thirty examples showing supercritical Hopf bifurcations. (b) Three examples of loss of self-sustained oscillations through other bifurcations.

**Figure S4**

Results of evolutionary calculations in which a target expression profile for gene 10 was assumed to depend on the daylength (*a, b*) or to be a gating type (*c*–*e*). (*a*) The target expression profile that was examined as an example of profiles, the forms of which depend on daylength. The profile has a peak at every dawn and a trough at every dusk. The mathematical form of the function was sin(*πt/τ*_L_) under light and sin(*π*[1 + (*t − τ*_L_)*/τ*_D_]) under dark where *τ*_L_ and *τ*_D_ represent the lengths of the light and dark periods, respectively. In the top, middle, and bottom panels, the light periods were set to 8 h, 12 h, and 16 h, respectively. Regular sinusoidal curves are shown as references (dotted). (*b*) Frequencies of gene regulatory networks (GRNs) showing self-sustained oscillation under at least one of the constant darkness or constant light exposure among 1,000 GRNs obtained after 1,000 generations of evolutionary calculation. The horizontal axis indicates daylength. (*c*) The target expression profiles that were examined as examples of gating type profiles. The mathematical form of the function is sin(2*π*(*t/τ*)^*p*^), where *τ* represents the lengths of light and dark periods. The Greater *p* resulted in a sharper peak and flatter trough. Note that the function is the same as a regular sinusoidal function when *p* = 1. (*d*) Frequencies of self-sustained oscillators among 1,000 GRNs obtained after 1,000 generations of evolutionary calculations. The horizontal axis indicates the value of parameter *p*. When *p* = 1 (i.e., for the regular sinusoidal curve), self-sustained oscillators were more frequently found in the environment with seasonality, in which a GRN was independently exposed to the 8L16D, 10L14D, 12L12D, 14L10D, and 16L8D environments. For *p* greater than 10^3*/*8^, they were more frequently found in the tropical environment, in which a GRN was exposed to only a 12L12D environment. (*e*) Distributions of cost values for the outcome GRNs from 1,000 different trials with different initial GRNs of evolutionary calculations with target profile *p* = 10. A cost value represents the deviation from the target profile. In conditions labelled as 12L12D, a GRN was exposed to a twelve-hour light period and a twelve-hour dark period repeatedly 50 times, and a cost value was calculated on the last 24-hour period. In conditions labelled as 8L16D– 16L8D, a GRN was independently exposed to 8L16D, 10L14D, 12L12D, 14L10D, and 16L8D environments, and cost values of those five environments were averaged. GRNs were classified according to behaviours under constant darkness and constant light exposure: purple for a GRN showing self-sustained oscillations under both constant environments, magenta and cyan for a GRN showing self-sustained oscillations only under light and dark conditions, respectively, dark grey if it does not show self-sustained or damped oscillations in any constant environment, and light grey otherwise.

