## Supplementary figures and images for "Evolution of self-sustained circadian rhythms requires seasonal change of daylight"

### Supplemental Figure 1

#### Figure S1

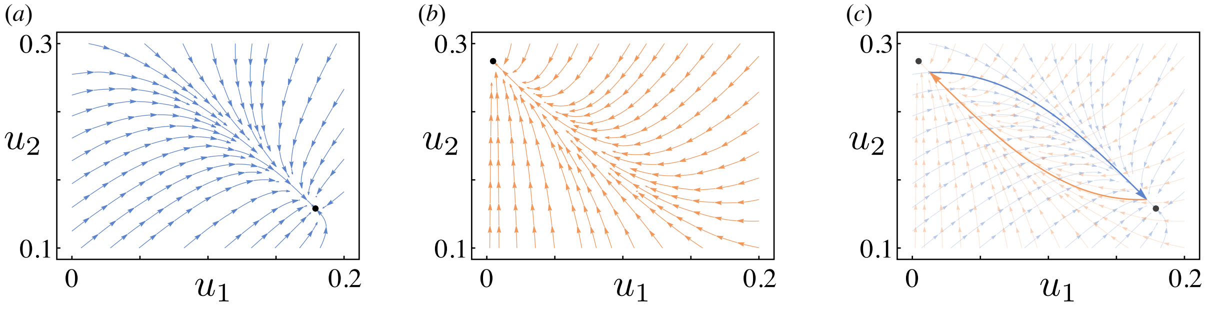

### Supplemental Figure 2

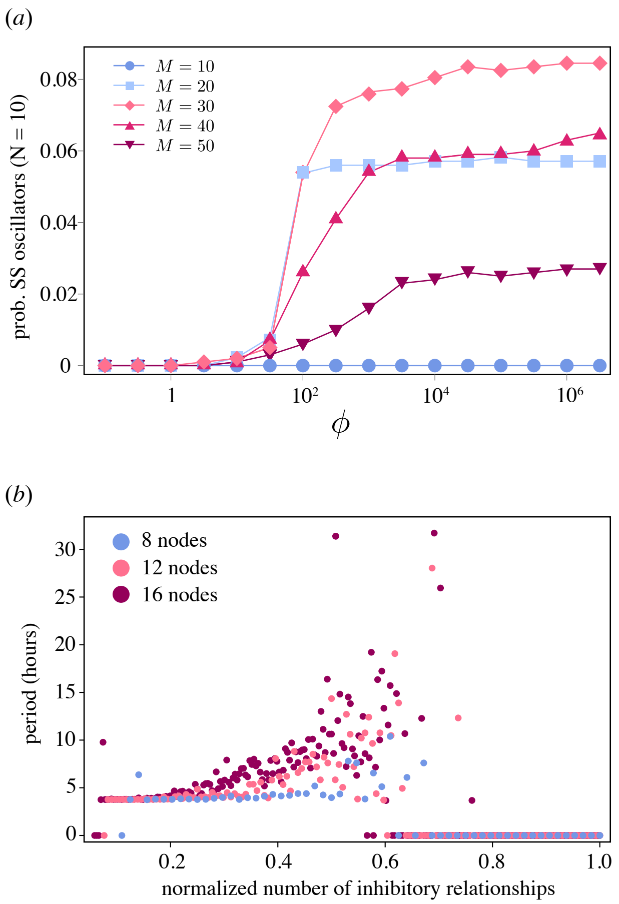

### Supplemental Figure 3

#### Figure S3

##### (a) Supercirtical Hopf bifurcation

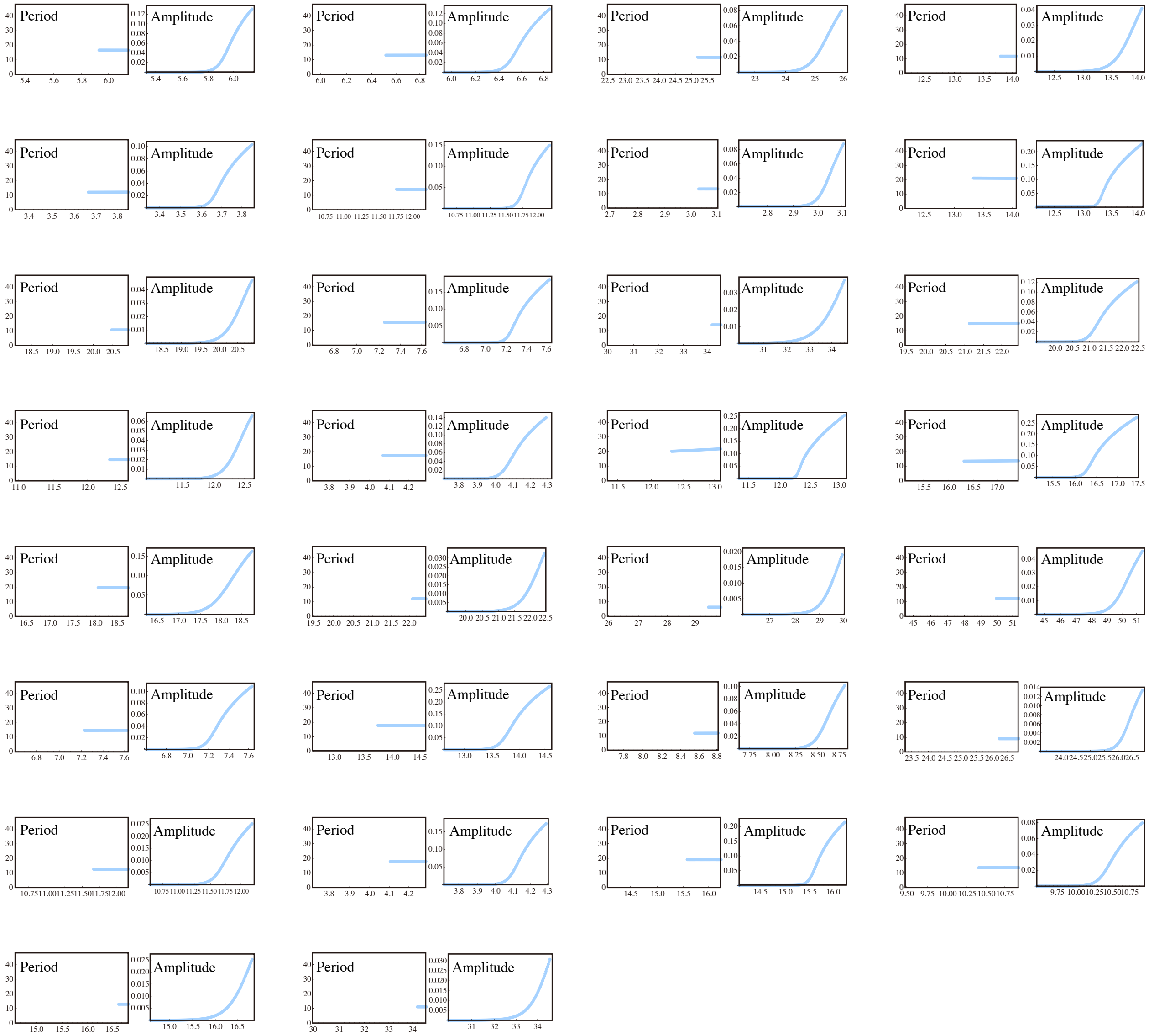


Time (h)

##### (b) The other birfucations

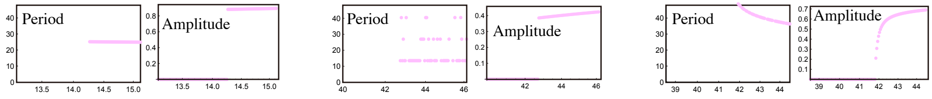


Time (h)

### Supplemental Figure 4

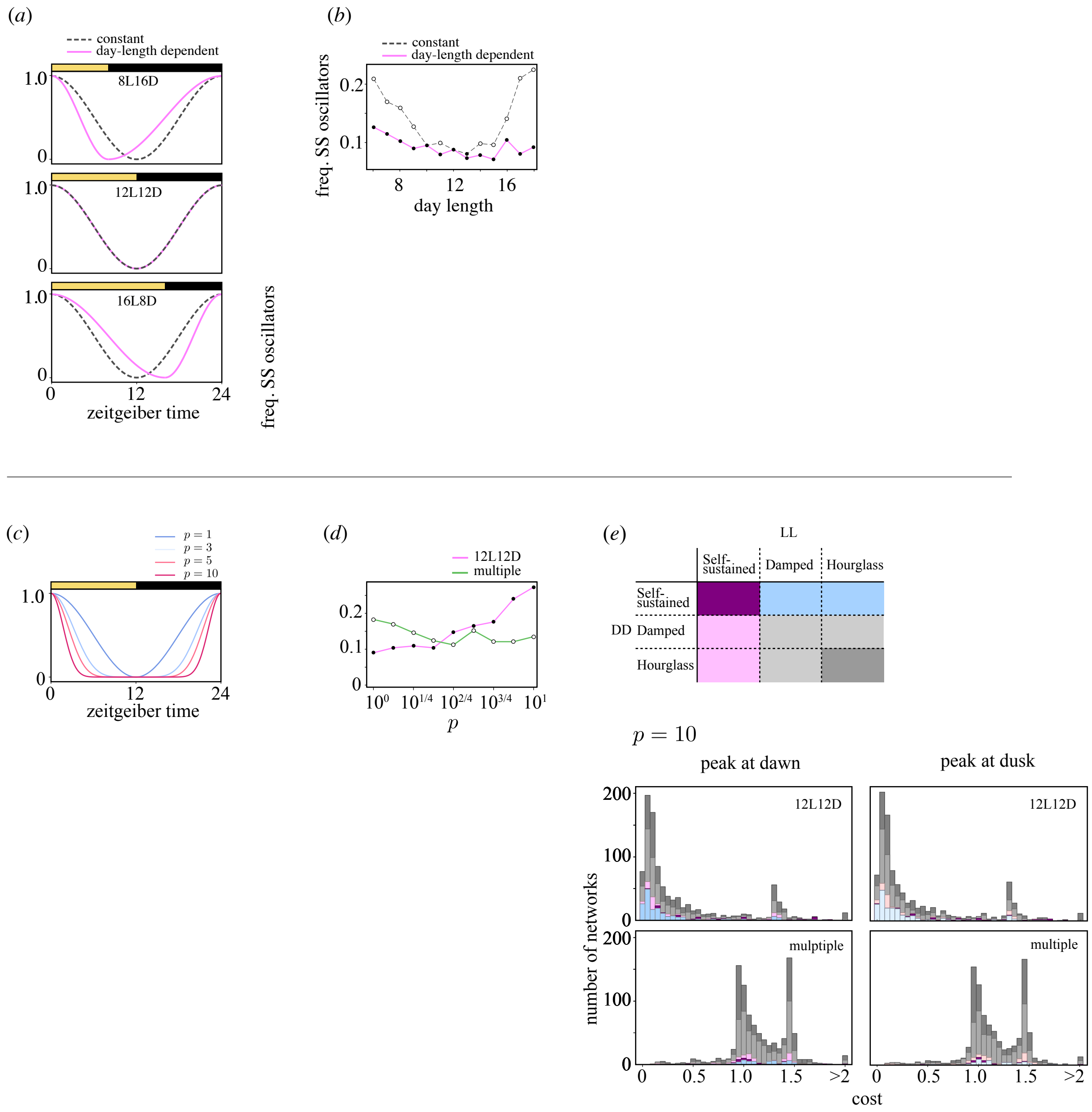
